# Label-free structural imaging of plant roots and microbes using third-harmonic generation microscopy

**DOI:** 10.1101/2024.04.13.589377

**Authors:** Daisong Pan, Jose A. Rivera, Max Miao, Peter Kim, Tomáš Tyml, Cristina Rodríguez, Umaima Afifa, Bing Wang, Yasuo Yoshikuni, Nathalie H. Elisabeth, Trent Northen, John P. Vogel, Na Ji

## Abstract

Root biology is pivotal in addressing global challenges including sustainable agriculture and climate change. However, roots have been relatively understudied among plant organs, partly due to the difficulties in imaging root structures in their natural environment. Here we used microfabricated ecosystems (EcoFABs) to establish growing environments with optical access and employed nonlinear multimodal microscopy of third-harmonic generation (THG) and three-photon fluorescence (3PF) to achieve label-free, *in situ* imaging of live roots and microbes at high spatiotemporal resolution. THG enabled us to observe key plant root structures including the vasculature, Casparian strips, dividing meristematic cells, and root cap cells, as well as subcellular features including nuclear envelopes, nucleoli, starch granules, and putative stress granules. THG from the cell walls of bacteria and fungi also provides label-free contrast for visualizing these microbes in the root rhizosphere. With simultaneously recorded 3PF fluorescence signal, we demonstrated our ability to investigate root-microbe interactions by achieving single-bacterium tracking and subcellular imaging of fungal spores and hyphae in the rhizosphere.

## INTRODUCTION

Root biology research informs wide-ranging areas of intensifying global interest including soil remediation^1^, climate^2^, genetics^3^, and sustainable agriculture^4,5^. Optical microscopy, capable of visualizing plant morphology at sub-cellular resolution, is critical for understanding plant structure and function^5^. However, compared to more extensively studied plant organs such as stems and flowers, a dearth of imaging studies on plant roots has resulted in roots being continually designated the “hidden half” of the plant body^6^.

The native environment of plant roots, composed of soil and organic matters, strongly scatters and absorbs light, severely limiting optical access to plant roots. As an alternative, microfabricated ecosystems such as EcoFABs^7^ provide controllable and reproducible growth conditions to plants. The absence of soil in EcoFABs relieves the requirement of uprooting the plant from its growing environment and provides non-destructive optical access for *in situ* imaging of root structure and its immediate microenvironment^8,9^.

Single-photon fluorescence microscopy techniques such as confocal microscopy^10^ and light sheet microscopy^11–13^ have been used to image plant roots^14,15^. However, light scattering by root tissues has limited their applications to relatively transparent or cleared root samples^16^. They also often require the introduction of extrinsic fluorescent labels. Transgenic fluorescent markers are compatible with live root imaging but can only be implemented for plants with established transformation systems and even in the best cases are laborious, whereas vital stains can suffer from poor incorporation^17^ or unwanted interference with cellular activity^18^. Thus, a label-free microscopy approach that can image structures at high resolution in opaque live root tissues would be highly desirable but has yet to be demonstrated.

Nonlinear optical microscopy methods utilizing near-infrared excitation light provide greater depth penetration in tissues than single-photon fluorescence techniques, and have been used to image both fluorescently labeled and unlabeled plant tissues^19–23^. Two nonlinear imaging modalities, three-photon fluorescence (3PF) microscopy and third-harmonic generation (THG) microscopy, have recently emerged as a powerful technique to image subcellular resolution at millimeter imaging depths in opaque tissues such as the mouse brain^24,25^. Here, we explored the potentials of using 3PF and THG microscopy for label-free *in situ* imaging at subcellular resolution of live plant roots in EcoFABs, including *Brachypodium distachyon*, a monocot grass known for its genetic tractability and relevance to agricultural crops^26,27^, and the dicot *Arabidopsis thaliana*, a central model organism in plant research^28^. We found that THG provided excellent optical resolution and label-free contrast for a variety of subcellular structures, while 3PF from the intrinsic fluorophores of root cells provided complementary structural information. We were able to visualize root hairs, vasculature, as well as subcellular components of mitotically active and border-like cells. In the optically opaque root of *B. distachyon*, THG microscopy imaged vasculature beyond 200 μm in depth in the mature zone and imaged through the entire 230-μm thickness of a root tip. Furthermore, we combined simultaneous label-free and fluorescence imaging to study root-microbe interactions, including real-time monitoring of *Pseudomonas simiae* dynamics in the vicinity of *A. thaliana* roots at single-bacterium resolution and visualizing the fungus *Trichoderma atroviride* adjacent to *B. distachyon* roots at subcellular resolution.

## RESULTS

### THG microscopy provides label-free contrast for plant root structures at subcellular resolution

THG, a coherent optical process that can convert three near-infrared excitation photons to one visible photon having one-third the excitation wavelength, has its signal originating from the third-order nonlinear susceptibility^29^. Under tight focusing conditions in microscopy, its contrast derives from the local optical heterogeneities within the excitation focus^19,30–32^. When the absorption of three near-infrared photons also promotes a fluorophore to its excited electronic state, 3PF can be generated by the same excitation laser. At longer wavelengths than the third-harmonic signal, 3PF signal can be spectrally separated from and simultaneously detected with the THG signal. The third-order nonlinear excitation involved in THG and 3PF confines the signal generation to within the excitation focus, thus optically sections 3D samples.

We conducted all imaging experiments using a custom-built multimodal microscope^25^ capable of simultaneously acquiring THG and 3PF signals (**Fig. 1a**). A near-infrared excitation laser beam (λ = 1300 nm; Monaco-Opera-F system, Coherent Inc.) propagated through a laser-scanning assembly comprising galvanometric mirrors (X,Y) and scan lenses to overfill a high-numerical-aperture (NA) water-dipping objective (Olympus XLPLN25XWMP2, NA 1.05, 25×). THG at 433 nm and 3PF (500 – 550 nm) signals were separated from the excitation light by a dichroic mirror (Dm) and further separated into two paths by an additional dichroic mirror for detection by photomultiplier tubes. We measured the point spread function (PSF) of our microscope by 3PF imaging of 0.2-μm-diameter fluorescent beads. The PSF had a lateral full width at half maximum (FWHM) of 0.65 μm in the xy plane and an axial FWHM of 1.78 μm (insets, **Fig. 1a**).

**Figure 1:**
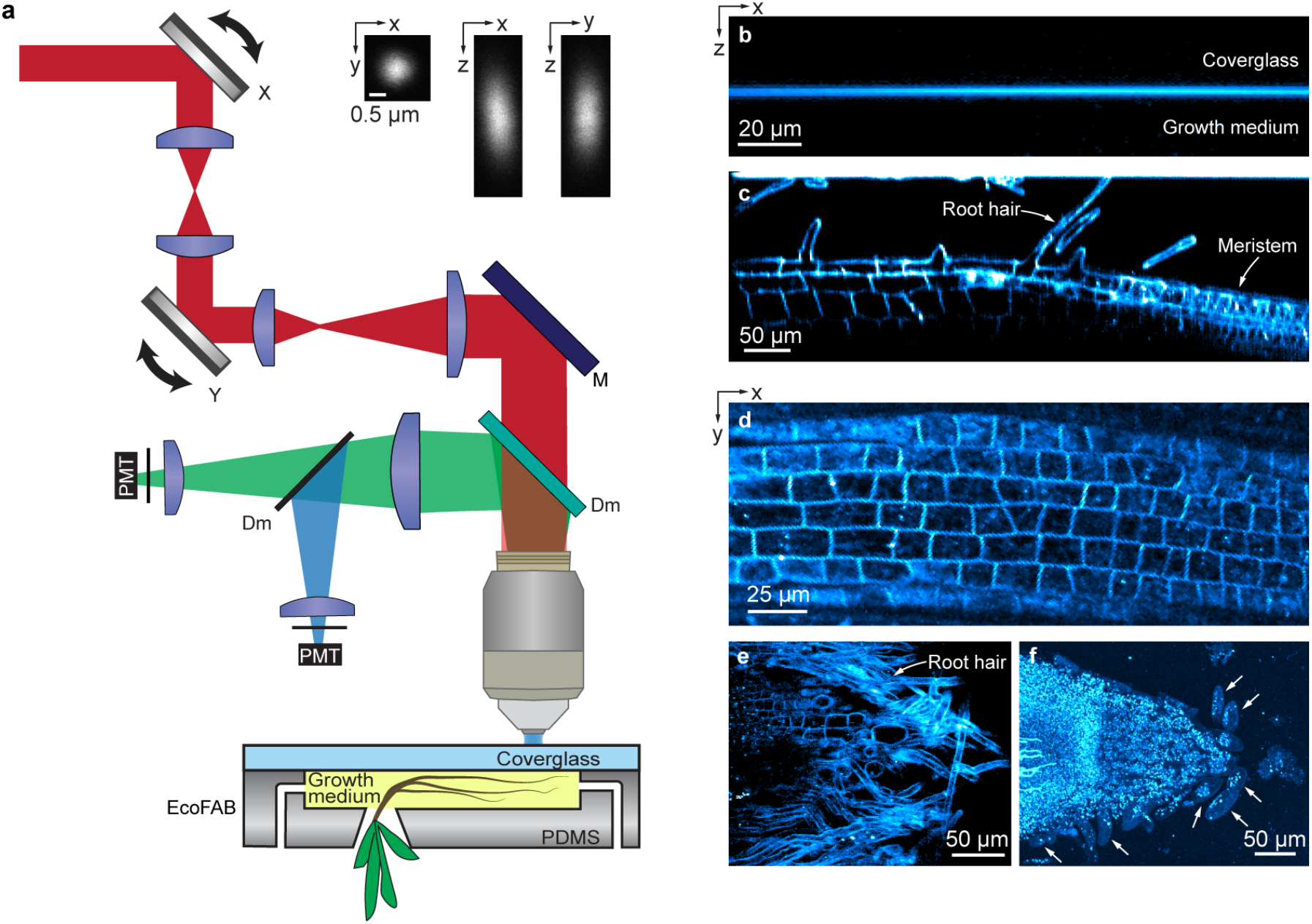
THG microscopy reveals plant root structure at subcellular resolution. (**a**) Schematics of the microscope and EcoFAB. Red: excitation light; Green: 3P fluorescence; blue: THG. X, Y: X and Y galvanometer mirrors; M: mirror; Dm: dichroic mirror; PMT: photomultiplier tube. Insets: lateral (xy) and axial (xz, yz) images of a 0.2-μm-diameter bead. (**b**) xz THG image of the interface between coverglass and growth medium. (**c**) xz and (**d,e**) xy THG images of *B. distachyon* roots inside the EcoFAB chamber. (b,c) acquired at 2 μm/pixel in X and 1 μm/pixel in Z. (d,e) acquired at 1 μm/pixel. (**f**) Brightest-spot projection with depth cueing (100% to 50%) of an 88-μm-thick image stack of a root tip acquired at 0.8 μm/pixel and z step size of 2.75 μm. White arrows: border-like cells. Post-objective power: (**b**) 2 mW, (**c,d**) 3 mW, (**e**) 4 mW, (**f**) 2.4-2.8 mW.

Seedlings of *B. distachyon* or *A. thaliana* were allowed to germinate within an EcoFAB growth chamber^9^ made of coverglass and PDMS before loading into the microscope (Methods). The excitation light entered through the coverglass side of the chamber and the emitted THG and 3PF signals were collected and detected in the epi direction. When the excitation focus was entirely within the coverglass or the growth medium, there was no THG signal generated due to the optical uniformity of the material within the focus^29,30^. When the excitation focus bisected the coverglass-medium interface, a strong THG signal was generated due to the abrupt change of susceptibility from glass to growth medium (**Fig. 1b**).

When the excitation focus was scanned across *B. distachyon* roots, THG signal provided high-resolution label-free visualization of various structures in both axial (**Fig. 1c**) and lateral (**Figs. 1d-f**) planes. Within the root, plant cells appeared as individual compartments (**Figs. 1c-e**), with strong THG signal observed at their cell walls, likely due to the different optical properties of the cell wall and the cytopolasm^33,34^. We confirmed that the THG signal arose from cell walls by acquiring 3PF and THG images from root samples stained with Auramine O, a fluorescent dye known to stain plant cell walls. Compared to unstained roots, fluorescent staining substantially reduced THG signal of the cell walls and severely disturbed intracellular structures (**Supplementary Fig. S1**), indicating that extrinsic fluorescent labelling led to unwanted perturbations on root structure.

In an example axial (xz) image (**Fig. 1c**), we observed elongated cells with root hairs protruding from the root surface – a well-known characteristic of the mature root zone^35^. Here from left to right, cells progressively diminished in size, corresponding to the transition into the meristematic zone. A similar transition was observed in a lateral (xy) image (**Fig. 1d**). Another lateral image section of a slightly inclined root revealed root hairs enveloping the root’s surface (**Fig. 1e**). Imaging the tip of a root, we observed cells that were detached from the primary body of the root near the root cap region (white arrows, **Fig. 1f**), reminiscent of border cells^36,37^ and suggesting that roots grown in the EcoFAB system resemble roots grown in soil. These label-free high-resolution images motivated us to systematically explore the cellular and subcellular features revealed by THG in *B. distachyon* roots, as detailed below.

### THG and 3P autofluorescence microscopy imaging of the mature zone of *B. distachyon* roots

In addition to THG, intrinsic contrast can arise from the autofluorescence of endogenous chromophores in biological specimens. Spectrally separating the THG and 3P autofluorescence signals, we simultaneously acquired THG and 3P autofluorescence images in the mature zone of *B. distachyon* roots (**Fig. 2; Supplementary Video 1**).

**Figure 2:**
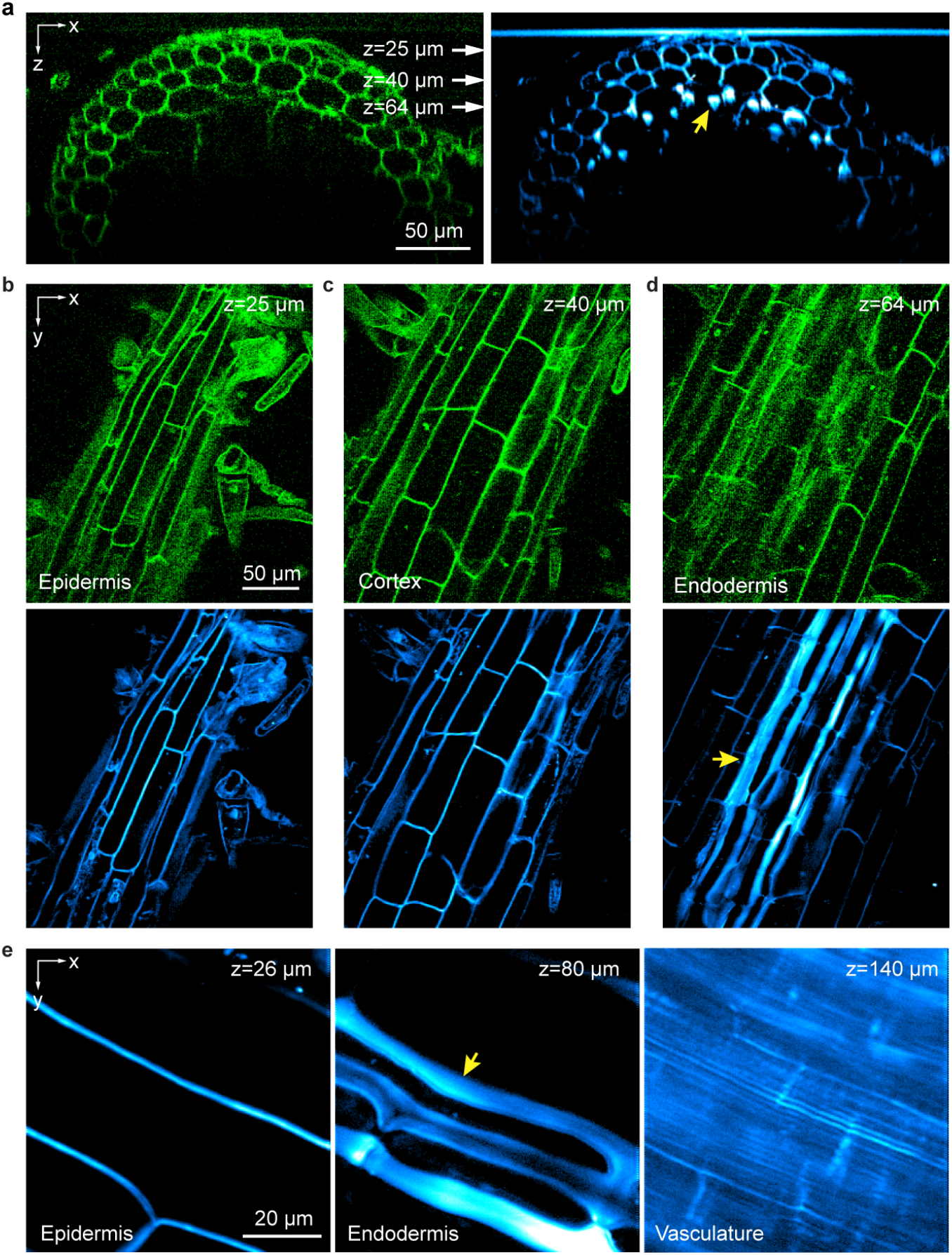
3P autofluorescence and THG microscopy visualize epidermis, cortex, endodermis, and vasculature in the mature root zone of *B. distachyon* roots. (**a**) 3P autofluorescence (green) and THG (cyan) xz images acquired at 1 μm/pixel showing a cross section of mature root zone. (**b-d**) xy images acquired at depths indicated by white arrows in **a** corresponding to (**b**) epidermis, (**c**) cortex and (**d**) endodermis tissues. Pixel size: 1 μm/pixel. (**e**) xy THG images of root tissues at 26 μm, 80 μm, and 140 μm depths, corresponding to epidermis, endodermis and vasculature, respectively. Pixel size: 0.3 μm/pixel. Post-objective powers: (**a-d**) 2.6 mW, (**e**) 1.2, 1.6, and 14 mW from left to right.

Roots in the mature zone contain concentric layers of epidermis, cortex, endodermis, and vasculature. In epidermis and cortex, THG and 3P autofluorescence signals co-localized at cell walls (**Fig. 2a-c**). Even though THG signal was ∼3.4–8.8× stronger than 3P autofluorescence signal, cell walls were easily visible in both channels. The contrasting composition of the plant cell wall and the surrounding cytoplasm generated THG signal, while the phenolic compounds in the cell wall^38–40^ were likely the source of autofluorescence.

Right before the THG and autofluorescence signals dropped off at larger depths, in the THG images we observed striation features of greater brightness and larger widths than the cell walls above (yellow arrows, **Fig. 2a,d,e**). Interestingly, the corresponding autofluorescence image did not exhibit this large increase of brightness (e.g., 65 – 83 µm, **Supplementary Video 1**), suggesting that THG signal was caused by a substantial change in the optical susceptibility. The location and morphology of these bright striated features of ∼10 µm width were consistent with Casparian strips in endodermis, which surround the vascular cylinder and regulate the passage of water and other solutes between cortex and vasculature^35^. The distinct molecular composition^41^ and thickness of Casparian strips from those of regular cell walls presumably led to its stronger THG contrast. Below the endodermis, vasculature structures were visualized as parallel channels ∼1.5 µm apart (**Fig. 2e**). Although ∼10× higher excitation power was needed for vasculature than for cells in epidermis and Casparian strips in endodermis, we were able to observe structures more than 200 µm deep into the mature root (**Supplementary Video 1**).

### THG and 3P autofluorescence microscopy image meristem and root tip of of *B. distachyon* roots at subcellular resolution

Compared with differentiated cells in the mature zone, cells in the meristem region contain more intracellular features required for root growth, which were visualized in the intracellular space in both 3P autofluorescence and THG cross-sectional images of a *B. distachyon* root (**Fig. 3a**). As in the mature zone, cell walls generated stronger THG signal than 3P autofluorescence signal and strong THG signal was observed in striated structures as those in the mature zone (white arrow, **Fig. 3a**).

**Figure 3:**
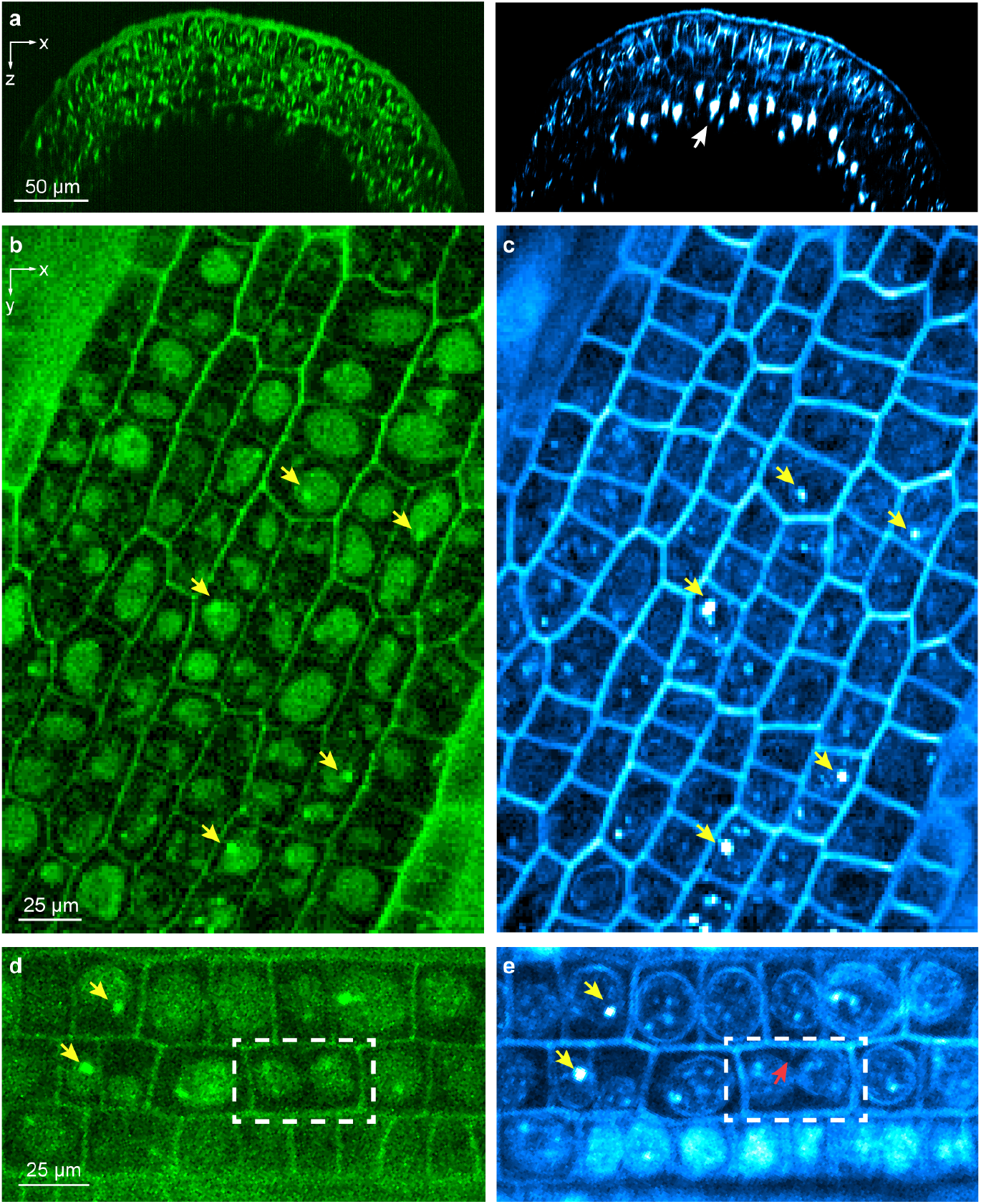
3P autofluorescence and THG microscopy provide label-free imaging of the *B. distachyon* root meristem at subcellular resolution. (**a**) 3P autofluorescence (green) and THG (cyan) xz images acquired at 0.3 μm/pixel showing a cross section of the root meristem. White arrow: putative Casparian strip. (**b-e**) 3P autofluorescence (green) and THG (cyan) xy images acquired at (**b,c**) 0.5 μm/pixel and (**d,e**) 0.3 μm/pixel. Yellow arrows: putative nucleoli; Dashed white box: a mitotic cell; red arrow: nascent cell wall. Post-objective power: (**a**) 5.3 mW; (**b,c**) 5 mW; (**d,e**) 7 mW.

In contrast to cells in the mature zone, cells in the meristem region were less elongated and approximately isodiametric (**Fig. 3b-e**). In 3P autofluorescence images, we observed ellipsoidal structures occupying most of the cell volume (**Fig. 3b,d**). There structures were likely enlarged nucleus, a characteristic feature of meristem cells, with their autofluorescence arising from the aromatic chemical structures of nucleic acid molecules themselves^42^. In several nuclei, we found bright and micron-sized autofluorescent aggregates (yellow arrows, **Fig. 3b,d**), which were consistent with nucleoli. The higher molecular density^43^ could explain the brighter fluorescence observed.

Due to the coherent nature and symmetry requirements of THG, subcellular structures exhibited distinct features in THG images (**Fig. 3c,d**) from those of 3P autofluorescence. When the excitation focus was within the optically uniform portion of the cell (e.g., cytoplasm or the non-nucleolus, nucleoplasm part of nucleus)^44^, THG signal was minimal. The varying susceptibilities across cytoplasm and nucleoplasm, however, gave rise to strong THG signal at the nuclear envelope (**Fig. 3e**). Inside the nuclei, strong THG signal was observed from the putative nucleoli (yellow arrows, **Fig. 3c,e**) due to their distinct optical susceptibility from nucleoplasm^44^. In both autofluorescence and THG images, nucleoli were often found near the nuclear envelopes, consistent with previous reports for plant cells^45,46^.

The distinctive features of meristem cell nuclei and the strong signal of cell walls in THG images provided us a label-free method to monitor cell division. For example, we observed a dividing cell with two daughter nuclei close to being separated (dashed box, **Fig. 3d,e**). In the equatorial plane^47^, a nascent cell wall could be detected in the THG image (red arrow, **Fig. 3e**).

THG also provided label-free structural contrast at subcellular resolution at the root tip encompassing the apical meristem and the root cap (**Fig. 4**). Imaging through a 230-μm-thick root tip (**Supplementary Video 2**), THG revealed a clear boundary between the meristem and the root cap^35^ (white arrows in **Fig. 4a**, a projected image of a 373 × 310 × 230 μm^3^ volume, and **Fig. 4b**, a single image section). Bright striated structures as in the endodermis image in **Fig. 2** were found within the THG images of the meristem region but not the root cap region (**Fig. 4a**; **Supplementary Video 2**), consistent with known anatomy of Casparian strips^35^. An axial cross-sectional view of the apical meristem region showed a complete encirclement of the central vasculature by the striated structures (**Fig. 4c**). Because light scattering and sample-induced aberration degraded focal intensity at deep depths, the Casparian strips in the bottom half of the root were substantially dimmer than those above. In cells protruding from the tip of the root cap, we observed bright granules of 3-7 μm in size (**Fig. 4d**), whose location and morphology were consistent with starch granules^48^. Throughout the apical meristem and root cap, we also saw smaller granules of 1-3 μm in size (**Fig. 4e**). We speculated that they may be processing bodies or stress granules^49^.

**Figure 4:**
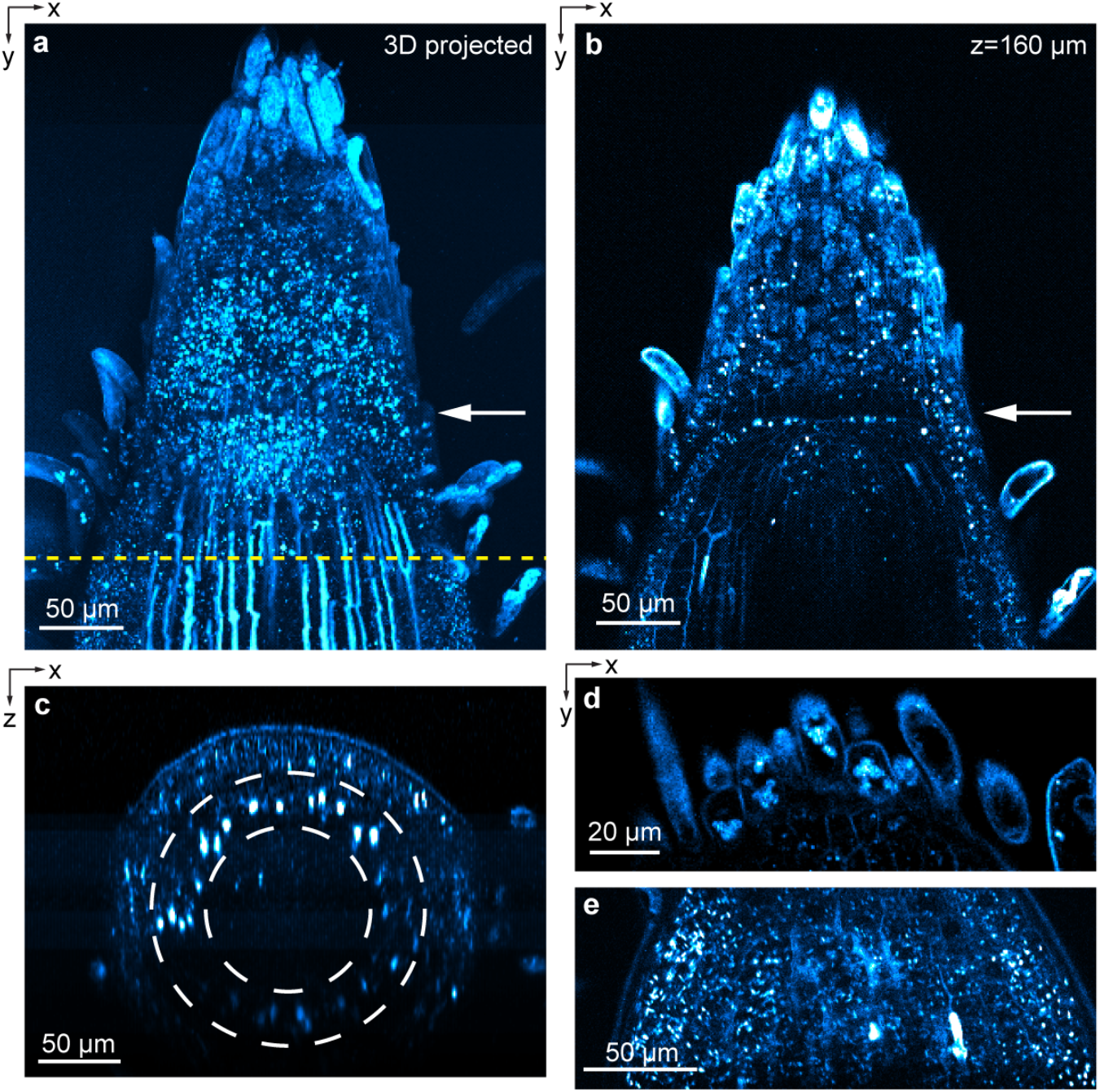
THG imaging of *B. distachyon* apical meristem and root cap. (**a**) Brightest-spot projection with depth cueing (100% to 50%) of an image stack through a root tip. 230-μm-thick image stack acquired at 1 μm/pixel and z step size of 2 μm. (**b**) xy image acquired at z=160 μm. White arrows in **a,b**: boundary between meristem and root cap. (**c**) xz images acquired along dashed yellow line in **a**. (**d,e**) xy images from two other root samples acquired at 0.5 μm/pixel. These roots were placed in between a microscope slide and a coverslip instead of inside EcoFAB to minimize sample motion. Post-objective power: (**a-c**) 3-15 mW; (**d**) 4mW; (**e**) 3.4 mW.

### THG and 3PF microscopy enable simultaneous imaging of plant roots and microbes in the rhizosphere

In addition to imaging plant roots themselves, the ability to simultaneously image microbes in the rhizosphere, the region in the vicinity of the roots where the microbiome interacts with the plant, would help understand the complex mechanisms through which root-microbe interactions impact plant growth^50,51^. We found that THG can be combined with 3P fluorescence to simultaneously image plant roots and bacteria as well as fungi in the rhizosphere *in situ*.

We first imaged two strains of *Pseudomonas simiae* bacteria^52^ near the surface of an *A. thaliana* root, including a GFP-labeled wildtype strain and a mutant strain without fluorescent labeling. Both *P. simiae* strains appeared as rod-like structures in THG images, often forming aggregates around the root tissue (**Supplementary Videos 3,4**; right panel, **Fig. 5a**). The GFP-labeled wildtype *P*.*simiae* had much stronger 3P fluorescence signal than the autofluorescence from root tissue (left panel, **Fig. 5a**) and showed up in both 3PF and THG channels at comparable signal strengths. During time-lapse imaging over 160 s at 1.1 Hz frame rate (**Supplementary Video 4, Fig. 5b**), we observed stationary (red and purple arrowheads), slowly moving (yellow arrowheads), as well as fast moving (orange arrowheads) bacteria near the *A. thaliana* root. These results indicate that our microscope is capable of simultaneously imaging and tracking of bacteria in the root rhizosphere, and that with additional fluorescence labeling for 3PF, it can image multiple bacterial strains simultaneously.

**Figure 5:**
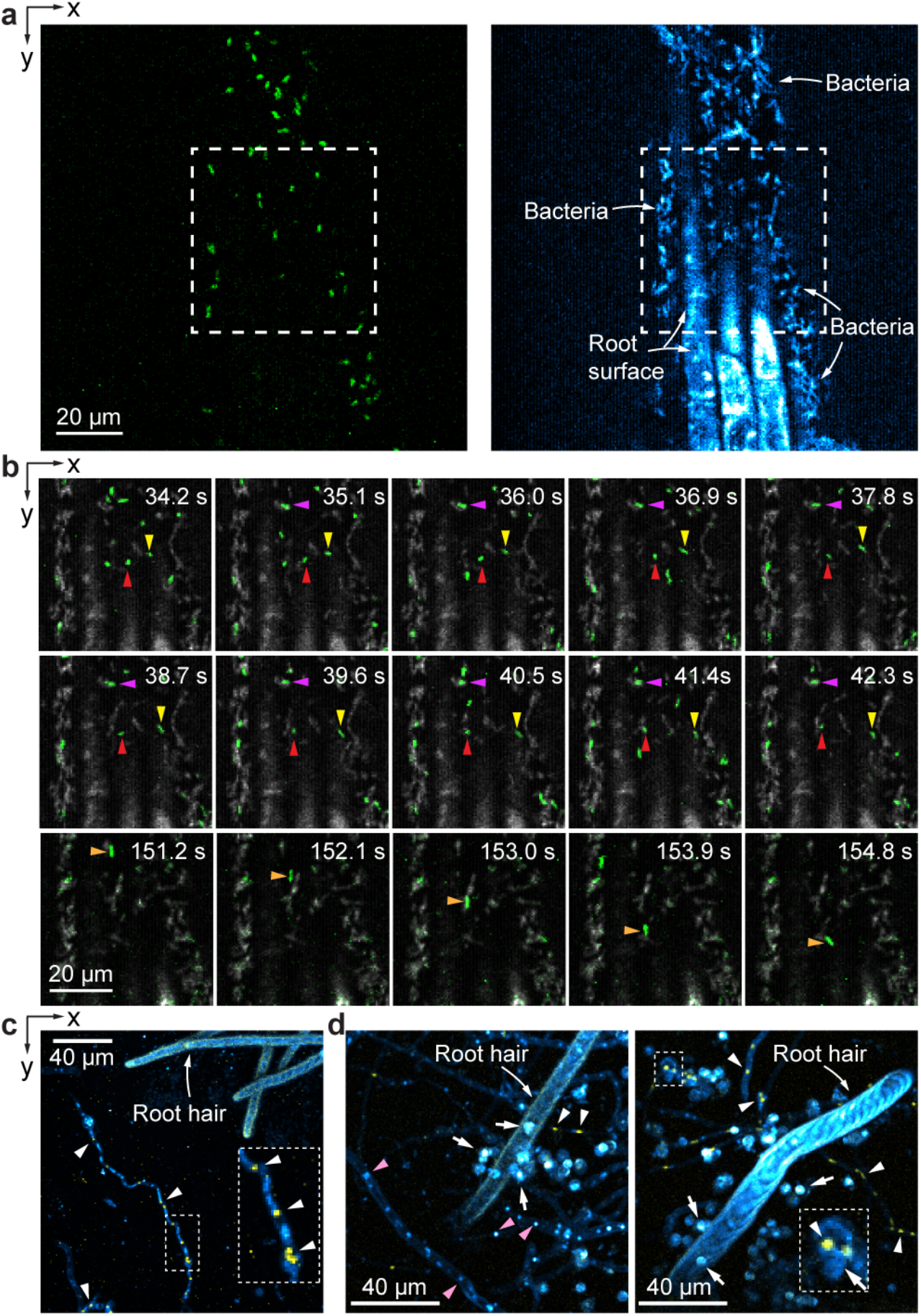
Imaging root-microbe interactions at high spatial and temporal resolution. (**a**) 3PF (green) and THG (cyan) xy images of *A. thaliana* root inoculated with two strains of *P. simiae* (wildtype *P. simiae* labeled with GFP, mutant *P. simiae* without GFP). (**b**) Consecutive frames of time-lapse imaging of the dashed box area in **a** with 3PF in green and THG in gray. Red/purple, yellow, and orange arrowheads: stationary, slowly-moving, fast-moving bacteria, respectively. (**c,d**) Maximal intensity projected 3PF (yellow) and THG (cyan) images of *B. distachyon* roots inoculated with *T. atroviride* strain IMI with GFP-labeled nuclei. (**c**) 50-μm-thick image stack acquired at 0.75 μm/pixel and z step size of 2.5 μm. (**d**) 70-μm-thick image stacks acquired at 0.5 μm/pixel and z step size of 2.5 μm. White arrowheads: GFP-labeled nuclei; pink arrowheads: unlabeled nuclei; white arrows: spores. Insets: zoomed-in views of white dashed boxes. Post-objective power: (**a, b**) 3.4 mW; (**c**) 5.6mW; (**d**) 5.3 mW.

We also investigated fungal colonization by imaging *B. distachyon* roots inoculated with *Trichoderma atroviride* strain IMI^53,54^, which had its nuclei labeled with GFP (H1-GFP^55^). Filamentous structures (**Fig. 5c**) with multiple fluorescent nuclei puncta of 1.5-2.5 µm in size^56^ (white arrowheads and inset, **Fig. 5c**) were identified as fungal hyphae. With their THG signal coming from fungal cell walls, these hyphae were observed near root hairs (**Fig. 5c**). In the THG image of another sample, we observed unlabeled punctate structures that are embedded within the hyphae and of similar size to the labeled nuclei, suggesting their identity as nuclei (pink arrowheads, **Fig. 5d**). In addition, spherical features with high THG signal were observed in close proximity to root hairs (white arrows, **Fig. 5d**). These spheres were 4-6 μm in diameter and were consistent with being fungal spores, with several of them having colocalized, GFP-labeled nuclei (inset, **Fig. 5d**). Therefore, THG microscopy proved to be a valuable tool for visualizing fungal hyphal structures, spores, and nuclei, alongside root structures. When combined with fluorescent labeling, the simultaneous detection of 3PF and THG signals can provide more specific structural insights into the interactions between roots and fungi.

## DISCUSSION

THG microscopy combined with microfabricated ecosystems allowed us to capture subcellular-resolution images of living plant roots without extrinsic fluorescent labels. Because THG signal originates from heterogeneity of optical susceptibilities within the excitation focal volume, it generates label-free visualization of cell walls. The 1.3-µm excitation light penetrated deep into the opaque tissues of *B. distachyon* roots, which are ∼2.5× thicker^57^ than the more widely studied and optically transparent roots of *A. thaliana*, and enabled us to visualize the vasculature in mature roots and image through the entirety of a 230-μm-thick root tip. Given that all cells in plant roots possess cell walls generating strong THG signal, THG microscopy can provide organ-scale views of root structures at subcellular resolution. In contrast to electron microscopy, which also offer a view of subcellular features, THG microscopy can be applied to live roots without labeling. Furthermore, due to its distinct wavelength, THG signal can be combined with simultaneously acquired fluorescent signals, either from the autofluorescence of endogenous molecules or from exogenous fluorescent labels, to provide structural context for biological processes of interest.

The structural features we observed in THG images are consistent with the known anatomy of plant roots. These include root hairs and elongated cells in the mature root zone. In both mature and meristem roots, we observed the layered arrangement of epidermis, cortex, and endodermis. Within endodermis, we found longitudinal striation features with strong THG signal that terminated at the root cap and were consistent with the location and morphology of Casparian strips. In the root meristem, THG contrast allowed us to visualize nucleoli and nuclear envelopes, providing information on stages of cell division. These structural identifications were strengthened by simultaneously recorded 3P autofluorescence signals from cell walls, nuclei, and nucleoli. In both apical meristem and root cap, between which a clear boundary can be identified in their THG images, the subcellular resolution of our imaging system allowed us to visualize and differentiate granules of varying sizes. Whereas starch granules were observed in root cap cells and border-like cells, we speculated that the small and bright puncta throughout meristem and root cap were likely stress-related granules, whose identities need to be further confirmed with molecular labeling approaches.

In addition to providing global structural information throughout the plant roots at subcellular resolution, THG microscopy also allows one to image bacteria and fungi in the rhizosphere. Because both bacteria and fungi have cell walls, they could also be visualized in a label-free manner by THG microscopy. Transgenic bacteria and fungi with fluorescent protein labels further improve the specificity of structural imaging. With multimodal THG and 3P fluorescence imaging, we were able to observe dynamics of bacterial distribution and fungal spores and hyphae near roots *in situ*. With deep penetration depth and optical sectioning capability, THG and 3P fluorescence microscopy therefore enable the investigation of root-microbe interactions throughout rhizosphere and within plant roots at high spatial and temporal resolution.

It should be noted, however, that plant cells are susceptible to light- and/or heat-induced damages, especially during multiphoton excitation^58,59^. In our experiment, extended imaging of meristem zone at post-objective powers over 10 mW always induced damage in the form of increasing the amount of bright puncta. Mature zone, on the other hand, withstood prolonged imaging at tens of mW without exhibiting visible damage. Care should always be taken to ensure that the biological process of interest is not unduly perturbed by imaging.

Our combined EcoFAB and multimodal imaging approach provides a powerful tool for studying the cellular structure of the roots. The large imaging depth of THG and 3P fluorescence microscopy enables the study of root-penetrating bacteria in opaque root tissues^60^. THG microscopy’s ability to visualize dividing root cells will enable studies on cellular division, elongation, and differentiation during root growth. Growth conditions could be altered within the EcoFAB chamber – providing a testbed for investigating how roots respond to environmental conditions, such as salinity or nutrient levels^61^, at high spatiotemporal resolution. In summary, by integrating microfabricated systems with nonlinear optical microscopy for label-free imaging of plant roots, we expect that our approach will illuminate the “hidden half” of the plant, shedding light on numerous unexplored facets of root biology.

## METHODS

### 3PF and THG microscopy setup

A simplified diagram of our multimodal (3PF and THG) microscopy is shown in **Fig. 1a**. The excitation source (not shown) consisted of an optical parametric amplifier (Opera-F, Coherent) pumped by a 40-W femtosecond laser (Monaco 1035-40-40, Coherent). Opera-F was tuned to generate 1,300 nm output at 1 MHz. A Pockels cell (M360-40, Conoptics) controlled the light power. A homebuilt single-prism compressor^62^ was used to cancel out the group delay dispersion (GDD) of the excitation beam path. The excitation laser beam was reflected by two conjugated galvanometric scanning mirrors (6215H, Cambridge Technology) and relayed to the back-pupil plane of a high NA water-dipping objective (Olympus XLPLN25XWMP2, NA 1.05, 25×) by two pairs of scan lenses (SL50-3P and SL50-3P, SL50-3P and TTL200MP; Thorlabs). The objective was mounted on a piezoelectric stage (P-725.4CD PIFOC, Physik Instrumente) for axial translation of the excitation focus. The fluorescence and THG signals were collected by the same objective, reflected by a dichroic mirror (FF665-Di02-25×36, Semrock) and detected by two photomultiplier tubes (H10770PA-40, Hamamatsu). An additional dichroic mirror (Dm, FF458-Di02-25×36, Semrock) and two filters (FF03-525/50-25 for fluorescence, FF01-433/24-25 for THG; Semrock) were used to split and filter the 3PF and THG signals. Frame rates were 0.2 – 0.6 Hz except for **Fig. 4**, which was acquired at 0.03 – 0.08 Hz, and **Fig. 5a,b**, which was acquired at 1.1 Hz.

### Bead sample

Carboxylate-modified fluorescent microspheres (FluoSpheres™, Invitrogen) were immobilized on poly(l-lysine)-coated microscope slides (12-550-12, Fisher Scientific).

### Imaging EcoFAB fabrication

Imaging EcoFAB devices were fabricated as described previously^9^. Negative molds for imaging EcoFAB were 3D printed using a Form2 printer (Formlabs) with clear resin version 4 (Formlabs). EcoFAB design can be obtained from https://eco-fab.org/device-design/. Each EcoFAB device was housed in a magenta box with a vented lid (MK5, with vented lid, Caisson Labs) for autoclave sterilization.

### *Brachypodium distachyon* growth conditions for EcoFAB imaging

*Brachypodium distachyon* line Bd21-3 seeds was used for this study^63^. Seeds were dehusked and surface sterilized in 70% ethanol for 30 s, followed by 50% v:v bleach (with 6.25% sodium hypochlorite chlorine) for 5 min, and rinsed 5 times with sterile milliQ water^9^. Seeds were then arranged on a sterile petri dish containing ½ Murashige and Skoog basal salt media (Caisson Labs) with 1% phytogel (Sigma-Aldrich). Surface sterilized seeds were stratified in the dark at 4 °C for 3 days. Following stratification, seeds were allowed to germinate in a growth chamber at 25 °C at 200 μmol·m^-2^·s^-1^, 16-hr light/8-hr dark. Three Days post germination, seedlings were transplanted into sterilized imaging EcoFABs containing 0.5× MS media with 0.8% phytogel^9^. Following transplantation, plants were grown in the growth chamber for three more days before imaging. All root samples were imaged within the EcoFABs except for **Fig. 4**, for which the plant was taken out of the EcoFAB and imaged with its root between two glass coverslips to reduce root tip motion.

### Dissected *Brachypodium distachyon* root sample

For dissected *B. distachyon* root samples imaged in **Supplementary Fig. S1**, staining was done using Bd21-3. *B. distachyon* seedlings 3 days post germination. Seedlings were dissected and fixed in 4% PFA in 1x PBS (Biotium) for at least 60 min with vacuum treatment. After fixation, seedlings were washed twice for 1 min in 1x PBS solution. For sectioned samples, primary root tissues were laterally hand sectioned using razor blades. Both hand sectioned and intact roots were then stained for lignin and suberin using 0.5% Auramine O solution (Sigma, CAS-No: 2465-27-2) adapted from Ursache et al., 2018^66^ before imaging.

### *Trichoderma* culture conditions and Inoculation

Three days post germination, sterile *B. distachyon* seedlings were inoculated with *Trichoderma atroviride* strain IMI^54^ containing nuclear GFP label (H1-GFP) (generously provided by Drs. Catherine Adams and Louis Glass, University of California Berkeley, CA, USA). Fungal spores were grown on PDA plates at 28 °C for 7 days 12/12 night/day cycle to induce sporulation. Spores were then harvested using sterile distilled water and separated from mycelia using a 0.4 micron filter (Pall). Spore concentration was determined using Neubauer chamber and then diluted to a spore suspension of 1×10^6^ spores/ml. Seedlings were soaked in the spore suspension for 2 hours prior to transplanting onto Imaging EcoFABs^64^. Following transplantation, plants continued growing in the growth chamber for two more days before imaging.

### *Arabidopsis thaliana* and *P. simiae* sample preparation

Seeds of *Arabidopsis thaliana* Col-0 (stock # CS66818) were obtained from the Arabidopsis Biological Resource Center (Ohio State University, Columbus, OH). Seeds were surface-sterilized by immersion in 70% (v/v) ethanol for 2 min, followed by immersion in 10% (v/v) household bleach containing 0.1% Triton X-100 (Roche Diagnostics GmbH) were stratified in distilled water at 4°C for 2 days. In this study, we utilized two strains of the root-colonizing bacterium *Pseudomonas simiae*, the non-fluorescently labeled strain *P. simiae* WCS417r and the eGFP-expressing strain *P. simiae* SB642, which has been previously characterized^52^. Both strains were pre-cultured under kanamycin selection (150 µg·ml^-1^) in Luria-Bertani medium (Sigma-Aldrich) diluted in 0.5× MS medium containing 2.15 g/L, 0.25 g/L of MES monohydrate (ChemCruz), and buffered to pH 5.7. The pre-cultured bacterial cells were washed twice with 0.5× MS medium and used to inoculate the stratified seeds of *A. thaliana* at an initial OD600 of 0.01 for each strain. The inoculated seeds were sown into agar-filled imaging EcoFAB chamber. The growth medium contained 0.5× Murashige and Skoog basal salt mixture (Sigma-Aldrich), 2.5 mM of MES monohydrate (ChemCruz), and was buffered to pH 5.7 and solidified with 1 wt% SFR agarose (Electron Microscopy Sciences). *Arabidopsis* seedlings were grown under 16 h light (140 µmol·m^-2^·s^-1^) and 8 h dark regime at 23 °C for 10-14 days.

### Digital image processing

Imaging data were processed with Fiji^65^. We used the ‘Green’ lookup table for 3PF images and the ‘Cyan hot’ lookup table for THG images. In **Fig. 5b**, the 3PF and THG signals were presented with ‘Green’ and ‘Gray’ lookup tables, respectively. For **Fig. 5c,d**, the 3PF and THG signals were presented with ‘Yellow’ and ‘Cyan hot’ lookup tables, respectively. To improve visibility, saturation and gamma of some images were adjusted. For **Fig. 5d**, we applied the ‘RemoveOutliers’ function in Fiji to eliminate hot pixels. We generated three-dimensional projection images (**Fig. 1f** and **Fig. 4a**) and a video (the second half of **Supplementary Video 2**) using Fiji’s ‘3D Project’ function with ‘brightest-point projection’ and depth cueing set at 100% to 50%.

## Acknowledgement

We thank Catherine Adams and Louis Glass at University of California Berkeley for providing *T. atroviride. B. distachyon* Bd21-3 seeds were obtained from Joint Genome Institute, Lawerence Berkeley National Labs (Berkeley, CA). The work conducted by the U.S. Department of Energy Joint Genome Institute (https://ror.org/04xm1d337), a DOE Office of Science User Facility, is supported by the Office of Science of the U.S. Department of Energy operated under Contract No. DE-AC02-05CH11231. This work was supported by the Lawrence Berkeley National Laboratory LDRD 20–116, LDRD 21-102, and Department of Energy DE-SC0021986.

## Author contribution

T.N., J.V., and N.J. conceived of the project; J.A.R. designed and built the microscope with help from C.R.; P.K. and M.M. prepared *B. distachyon* samples; D.P., J.A.R., P.K., and M.M. acquired *B. distachyon* images; M.M. prepared *B. distachyon* and *T. atroviride* samples; D.P. and M.M. acquired *B. distachyon* and *T. atroviride* images; T.T. and N.H.E. prepared *A. thaliana and P. simiae* samples; D.P. and T.T. acquired *A. thaliana and P. simiae* images; D.P., J.A.R. and N.J. wrote the manuscript with inputs from all authors.

## Data availability

All data generated or analyzed during this study are included in this published article and its Supplementary Information files.

## Notes

### Competing Interest Statement

The authors have declared no competing interest.

### Summary of Updates

Author list updated; supplemental figure added.

