## Supplementary Fig. S1 for "Label-free structural imaging of plant roots and microbes using third-harmonic generation microscopy"

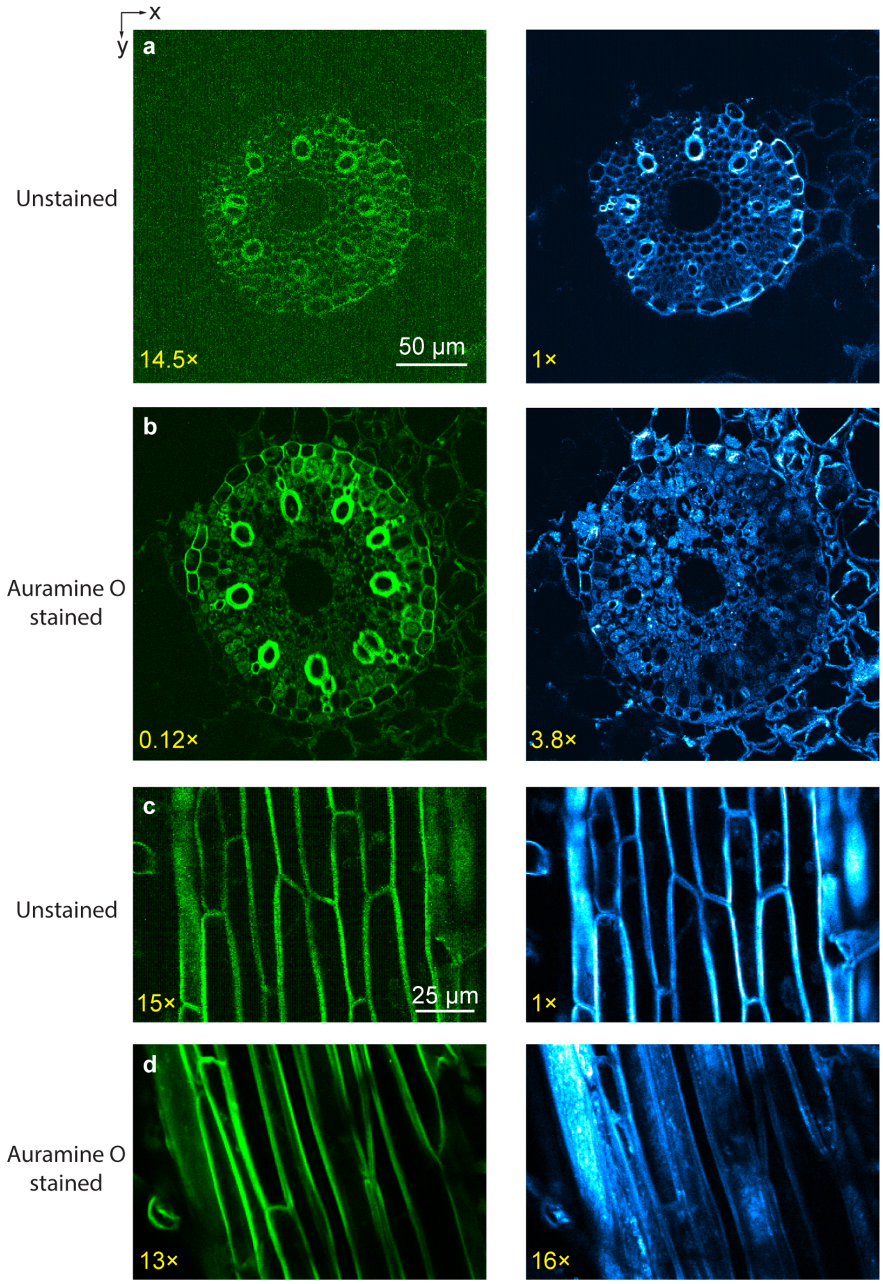


**Supplementary Figure S1:** Fluorescence staining disrupts the inner structures of root samples. (**a,b**) 3PF (green) and THG (cyan) xy images of transverse root sections of mature zones. (**c,d**) 3PF (green) and THG (cyan) xy images of lateral sections of mature zones from two root samples. Roots are (**a,c**) unstained or (**b,d**) stained with Auramine O to label lignin-rich structures, including cell walls. All images were acquired at 0.5 μm/pixel. The brightness of both 3PF and THG channels in (**a,b**) and (**c,d**) was normalized to their respective THG images of unstained roots, with digital gain values listed at the bottom left of each image. (Larger gains indicate dimmer images.) Clearly, staining reduces brightness in the THG channels in (**b,d**), indicative of structural disruption and decreased optical inhomogeneity within the cells caused by staining.
